# VECTOR: A Framework to Identify and Quantify Structural Changes in Chromatin using Hi-C Data

**DOI:** 10.1101/2025.09.24.678266

**Authors:** Kavana Priyadarshini Keshava, Dieter W. Heermann, Arnab Bhattacherjee

**Author notes:** Contributing authors.

## Abstract

Hi-C and related 3C technologies capture 3D genome architecture by measuring contact frequencies between genomic loci. These contact maps are often sparse and noisy, making comparison across samples challenging. Traditional metrics such as Pearson’s correlation fail to capture their structural complexity. To address this, we present VECTOR (Von nEumann entropy deteCTiOn of stRuctural patterns), a novel framework that uses Von Neumann entropy and graph spectral theory to quantify chromatin organization from Hi-C data. By constructing normalized graph Laplacians from contact matrices, VECTOR captures both short- and long-range chromatin interactions, offering a robust measure of 3D genome architecture. Applied to human and mouse datasets across diverse cell types, developmental stages, and replicates, VECTOR reveals reproducible entropy patterns that distinguish biological conditions, including cancer versus stem cells and early versus late embryonic stages. VECTOR outperforms existing Hi-C comparison tools such as ENT3C and HiCRep, especially in sparse datasets, by reliably differentiating biological replicates from non-biological ones. It further demonstrates stability across Hi-C resolutions and successfully detects genome-wide structural transitions linked to regulatory elements such as TADs and CTCF sites. When applied to sparse single-nucleus Hi-C data from mouse brain development, VECTOR accurately resolves developmental progression that is poorly captured by correlation-based methods. Our results establish VECTOR as a robust and interpretable tool for comparative Hi-C analysis, with broad applications in chromatin biology, development, and disease.

## 1 Introduction

The three-dimensional (3D) organization of the genome plays a fundamental role in gene regulation, chromatin architecture, and various cellular processes. Recent advances in chromosome conformation capture (3C)-based techniques, particularly Hi-C, have provided genome-wide insight into chromatin interactions [1–3]. These methods leverage proximity ligation followed by deep sequencing to map interactions between chromatin regions, uncovering key structural features such as chromatin loops, topologically associating domains (TADs), compartmental domains, and A/B compartments [2, 4, 5]. These higher-order chromatin structures are critical for coordinating gene expression and maintaining genome stability. One of Hi-C’s key advantages is its ability to detect long-range chromatin interactions, where regions that are distant in linear genomic sequence are spatially close in the nucleus, enabling regulatory mechanisms such as enhancer-promoter interactions [6–8].

Hi-C data are typically represented as contact matrices, where the genome is divided into fix-sized bins, and the matrix entries reflect interaction frequencies between different genomic loci. However, these matrices are often sparse and susceptible to technical noise arising from sequencing biases, incomplete sampling, and spurious ligation events [9]. Additional sources of noise include artifacts introduced by chromatin fragmentation and ligation protocols, which can obscure biologically meaningful interactions. Furthermore, sequence composition and mappability introduce systematic biases that complicate downstream analysis [10, 11]. To mitigate these challenges, similarity metric, such as Pearson’s correlation matrices, are frequently employed to reduce random fluctuations and enhance structural signal detection.

Despite technological advancements, distinguishing genuine chromatin interactions from experimental artifacts remains a significant challenge. Improving the accuracy and reproducibility of Hi-C data analysis requires the development of robust statistical frameworks and higher-resolution methodologies. Additionally, the computational demands of analyzing large Hi-C datasets present another hurdle, necessitating efficient algorithms and scalable analytical tools [12]. Integrating Hi-C data with complementary genomic datasets such as ChIP-seq and RNA-seq is essential for a comprehensive understanding of chromatin dynamics and gene regulation [13]; however, developing sophisticated methods to achieve this integration remains a challenge [11, 14].

A crucial aspect of Hi-C analysis is the comparison of datasets to investigate chromatin organization under different biological conditions. Various computational methods have been developed for this purpose. Correlation-based metrics, such as Pearson and Spearman coefficients, offer a broad measure of similarity between data sets but may not capture local structural nuances [9].Structural entropy has been explored as an effective approach to segment Hi-C contact maps into meaningful domains such as TAD. deDoc offers a fast, normalization-free method to detect TAD-like domains and determine optimal bin sizes, demonstrating robustness on pooled single-cell data [15]. SuperTAD formulates domain detection as a structural information minimization problem, solved efficiently via dynamic programming [16]. These entropy-based methods enable hierarchical multiscale analysis with minimal pre-processing or parameter tuning. However, deep learning-based methods are often less accessible, and domain-based representations like TADs may not be well-suited for distinguishing biological replicates or subtle structural variations. In contrast, distance-based methods [17] and the structural similarity index (SSIM) [18, 19]enable more detailed, region-specific comparisons. However, these approaches can be computationally intensive, particularly when incorporating deep learning techniques and polymer physics-based modeling [13, 20].

Alternative strategies include matrix-based methods such as the Stratum Adjusted Correlation Coefficient (SCC), which accounts for distance-dependent contact decay as in HiCRep [10], and network-based approaches that leverage graph theory to characterize chromatin interactions. Other advanced methods have been introduced to improve Hi-C dataset comparisons. For example, Selfish [21] quantifies local self-similarity, GenomeDISCO [22] employs random walks to smooth contact matrices before computing similarity scores, and HiC-spector [23] utilizes spectral graph theory to assess structural differences. Additionally, clustering algorithms and graph-based methods can reveal hidden patterns in Hi-C data by identifying chromatin domains, interaction hubs, and community structures. Techniques such as diffusion maps facilitate the identification of non-obvious structural patterns, while QuASAR-Rep and QuASAR-QC [24] focus on quality assessment and reproducibility, ensuring that Hi-C datasets are robust and reliable. The latest ENT3C similarity measure uses Pearson’s correlation of diagonal submatrix scanning with Von Neumann entropy [25].

Understanding the structural complexity of biological networks requires analytical tools capable of capturing both global organization and fine-scale connectivity patterns. Traditional graph-theoretical methods often rely on predefined topological descriptors, which may overlook subtle or distributed structural variations. In recent years, approaches grounded in quantum information theory have emerged as powerful alternatives, offering information-theoretic measures that account for the entire network structure. A key development in this direction is the use of *von-Neumann entropy*, a quantum analogue of Shannon entropy, which has been successfully applied to characterize complex systems such as brain functional networks [26–30]. This measure treats the network as a quantum-like system by associating it with a normalized graph Laplacian that satisfies the properties of a quantum density matrix. The resulting entropy reflects the degree of structural disorder in the network, integrating contributions from all eigenmodes of the Laplacian spectrum. As such, von-Neumann entropy provides a comprehensive, scale-resolved description of network architecture.

In this study, we extend this formalism to the domain of 3D genome organization. Specifically, we introduce VECTOR (*Von nEumann entropy deteCTiOn of stRuctural patterns in Hi-C data*), a computational approach for analyzing Hi-C contact matrices using von-Neumann entropy [25, 31]. VECTOR transforms each Hi-C contact matrix into a graph representation, applies a Laplacian transformation, and computes a one-dimensional entropy signal based on the eigenvalues of the normalized Laplacian. This signal captures both local and global features of chromatin structure.

While numerous techniques exist for comparing Hi-C data, no single method can comprehensively capture all structural variations due to the multiscale and context-dependent nature of chromatin organization. VECTOR addresses this gap by providing a scalable, model-agnostic, and information-theoretic framework for quantifying chromatin structure. Through its application, we demonstrate the utility of spectral entropy in capturing biologically meaningful patterns in 3D genome organization. We show that the similarity score of VECTOR remains highly robust to variations in sequencing depth and binning resolution, enabling precise differentiation between Hi-C contact matrices from different cell lines while reliably identifying biological replicates of the same cell line. Furthermore, we emphasize the potential of entropy signals in investigating both intra- and intercellular differences in 3D chromatin organization. By representing Hi-C data as a graph, VECTOR captures key structural features, providing valuable insights into chromatin architecture through entropy-based analysis. This metric quantitatively assesses structural variations across the genome, facilitating the differentiation of Hi-C matrices under diverse biological conditions, such as cell differentiation. With its computational efficiency and ease of implementation, VECTOR serves as a powerful tool to study the organization of 3D genomes.

## 2 Methods

### 2.1 Von Neumann entropy for weighted graphs

Hi-C data capture the spatial proximity of genomic loci, often revealing long-range interactions between distant genomic segments. The data are typically represented in a three-column format, where the first two columns denote the genomic coordinates of interacting loci, and the third column records the interaction frequency.

To quantify the structural organization encoded in these interactions, we employ the VECTOR framework, which models Hi-C contact matrices as weighted graphs. For a given *N* × *N* contact matrix, VECTOR treats each genomic locus—binned at fixed resolution as a node, with edge weights corresponding to the interaction frequencies between pairs of loci. This graph-based representation enables downstream analysis of chromatin architecture using tools from network theory and spectral graph analysis.

To quantify the complexity of these graphs, we use the *von-Neumann entropy*, a graph-theoretic measure rooted in quantum information theory. This entropy generalizes classical concepts of entropy to networks, allowing us to characterize the structural disorder and connectivity in weighted graphs.

#### Graph Construction from Hi-C Data

Let *G* = (*V, E*) denote a weighted graph, where *V* is the set of nodes (genomic bins) and *E* is the set of edges representing contact frequencies. The adjacency matrix *A* is defined by:

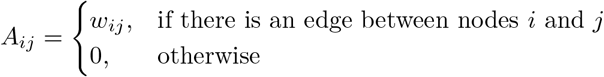

where *w*_*ij*_ is the contact frequency (edge weight) between nodes *i* and *j*. Next, we compute the degree matrix *D*, a diagonal matrix where each element *D*_*ii*_ is the sum of weights incident to node *i*:

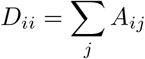

Using *A* and *D*, we define the combinatorial Laplacian:

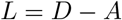

From this, we compute the normalized Laplacian matrix:

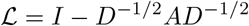

where *I* is the identity matrix and *D*^−1*/*2^ is the inverse square root of the degree matrix.

#### Von Neumann Entropy Calculation

Let *λ*_*i*_ be the eigenvalues of the normalized Laplacian matrix ℒ. These eigenvalues encode the structural characteristics of the graph. To compute the von-Neumann entropy, we first normalize the non-zero eigenvalues to form a probability distribution:

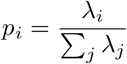

The von-Neumann entropy of the graph is then defined as:

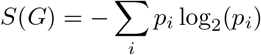

To capture local structural variation along the genome, entropy is not computed over the entire contact matrix but rather on localized ego-graphs centered on sampled nodes. Specifically, for a given stride parameter *n*, VECTOR selects every *n*^th^ node as a center and constructs a 1-hop ego-graph consisting of the node and all its immediate neighbors based on contact frequency. For each ego-graph, the corresponding adjacency and Laplacian matrices are constructed, and von-Neumann entropy is computed as described above. To quantify the complexity of graph structures derived from Hi-C contact matrices, entropy values are computed and scaled to the [0,1] range using Min-Max normalization. These entropy values often vary across genomic bins, with some exhibiting unusually low interaction frequencies or connecting only to adjacent nodes. Such sparsely connected bins result in significantly reduced entropy, often reflecting experimental noise or structural artifacts.

To identify and remove these anomalous bins, we apply the 1.5× interquartile range (IQR) rule. Bins with entropy values below Q1-1.5xIQR or above Q3+1.5xIQR are flagged as outliers and excluded from downstream analysis. This approach ensures robust filtering of extreme values without making distributional assumptions.

Subsequently, the remaining entropy values are rescaled using Min-Max normalization, allowing for consistent comparison across samples and experimental conditions while minimizing the influence of outliers (Detailed Workflow Supplementary Fig 4).

#### Similarity Score

When comparing two contact matrices, VECTOR measures similarity using the Pearson correlation coefficient between the entropy signals between them . Values are excluded from the calculation if they are missing in any of the matrices being compared.

#### VECTOR Parameters

In VECTOR, stride refers to the step size used to sample nodes from an N×N contact matrix to generate local ego-graphs for entropy computation. For a stride n, every nth node is selected (e.g., node 0, node n, node 2n, etc.). For each selected node, an ego graph is extracted - typically including the node and its immediate neighborhood - based on the contact matrix. The von-Neumann entropy is then computed (see Supplementary Fig 4) for each of these ego-graphs to form the entropy signal over the genome.

#### Cell Lines and their Sources

##### Hi-C Datasets

Processed Hi-C data for the A549, G401, and LNCaP cell lines were obtained from the ENCODE Portal (ENCODE Project, last accessed November 2024). We downloaded BAM files (ENCODE4 v1.10.0, GRCh38) for each replicate and converted them into pairs files using the pairtools parse function (see Supplementary Table1 and Table2). These datasets were selected for comparability with a recent review on contact matrix similarity metrics.

The datasets include established cell lines and various developmental stages of mouse embryonic cells, as well as human cancer and non-cancer cell lines. Mouse embryonic stem cells (mES) originate from the inner cell mass (ICM) of the blastocyst and serve as a key model for studying early development. The PN3 and PN5 stages represent crucial zygotic reprogramming events, while the Early 2-cell, Late 2-cell, and 8-cell stages mark progressive genome activation and chromatin reorganization. The Cortex was included to compare embryonic transitions with differentiated neuronal tissue.

Human datasets include hESCs, pluripotent cells used in regenerative medicine. Cancer cell lines analyzed include A549 (lung carcinoma), LNCaP (prostate cancer), and G401 (Wilms’ tumor), chosen for their relevance in disease research. By studying these diverse cell types, we aim to capture key chromatin and contact matrix changes during development and disease progression. Additionally, single-cell Hi-C data from mouse embryos were incorporated to assess VECTOR’s robustness and reproducibility in sparse matrix conditions. Details of these input datasets are summarized in Supplementary Table 3.

##### Binning and Contact Matrix Generation

Using Juicer Tools `Juicer tools (juicer_tools_1.22.01.jar)` [32], `pairs.gz` files were binned into contact matrices and stored in `.txt` format.

##### Down-sampling pairs

When specified, pairs files were down-sampled using the pairtool sample function (pairtools, version 0.3.0) [33].

##### CTCF Sites and TADs Detection

Mouse CTCF binding sites were obtained from the CTCF database version 2.0 [34, 35]. Topologically associating domains (TADs) were identified using the Arrowhead algorithm implemented in the Juicer pipeline, with a window size of 2000 bins at a resolution of 50 kb.

##### Contact Matrix Balancing

VECTOR can operate on balanced or unbalanced contact matrices. Unless stated otherwise, all tests were performed on raw interaction frequency matrices.

##### Identifying Similar Regions Between Two Contact Matrices Using Entropy Signals

To identify structurally similar regions between two genomic contact matrices, we calculate entropy values across chromosome 2 (from 60 to 110 MB). These entropy values represent the complexity of genomic interactions. We then compare the entropy profiles of the two matrices using linear regression. Regions with entropy values that have the smallest perpendicular distance to the fitted regression line indicate similar structural complexity. From this, we select the top 10 regions with the smallest deviations from the regression fit and identified genes curated by NCBI within these regions using the UCSC browser [36].

## 3 Results

### 3.1 Tracking 3D genome changes through VECTOR entropy signals

Hi-C data capture the spatial interactions between linearly distant genomic segments. We utilized the graph-based approach VECTOR to quantify this organization. Given an N×N contact matrix, VECTOR models it as a graph, where nodes represent genomic locations (in base pair bins) and the edges correspond to interaction frequencies. The entropy is then calculated using the von-Neumann entropy of eigenvalues derived from the normalized Laplacian matrix.(Fig. 1A and Section 2).

**Fig. 1.**
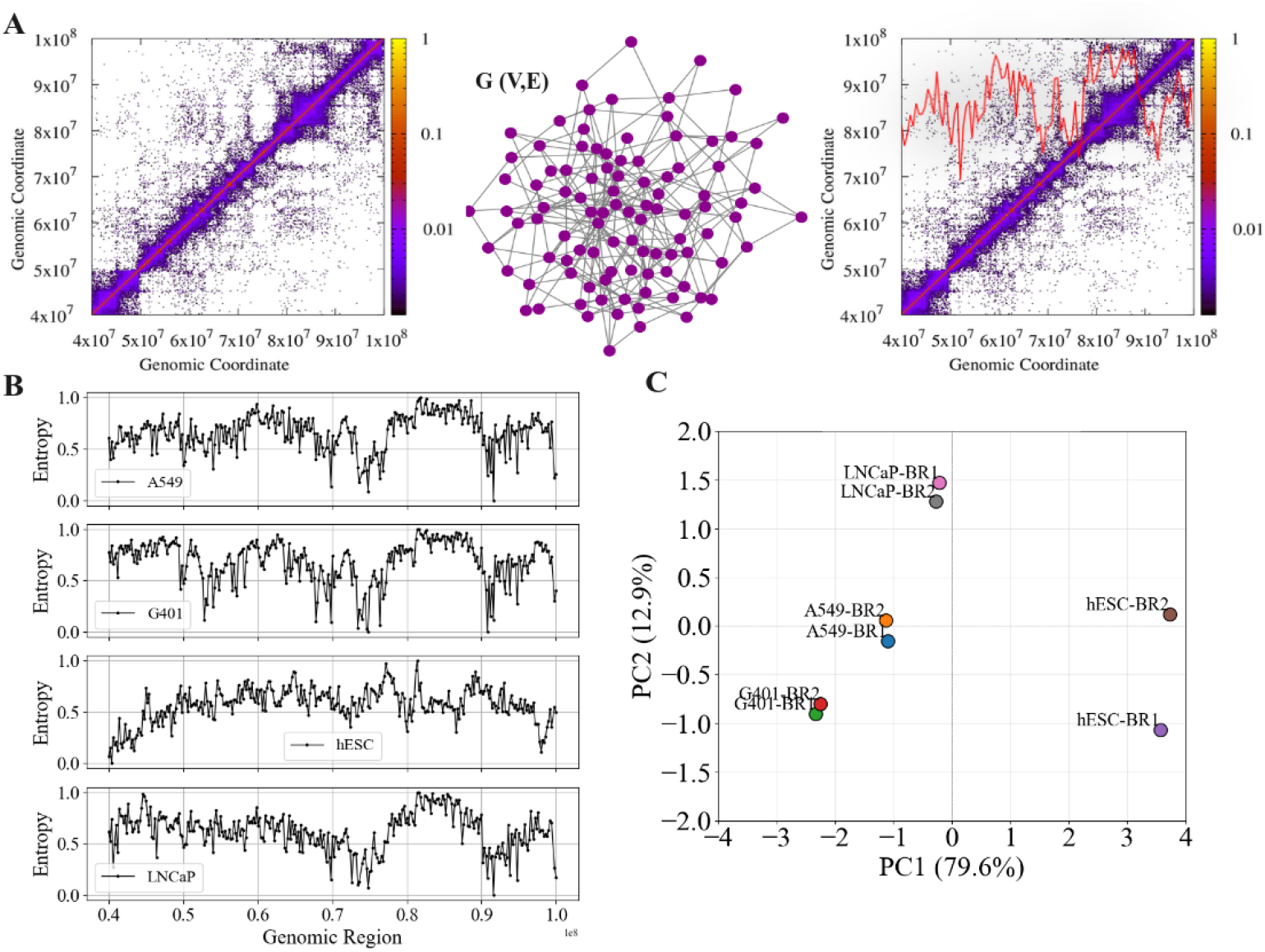
(A) Schematic representation of the entropy method, illustrating how VECTOR derives entropy signals using the graphical von-Neumann approach applied to an input contact matrix. (B) Entropy values across chromosome 14 for different cell lines. (C) PCA plot of entropy values from contact matrices of four human cell lines for chromosome 14(region 40e+07 -100e+07), demonstrating strong clustering along PC1. The most outlying sample corresponds to biological replicate 2 of hESC.

As a test case, we analyze the entropy signal of human chromosome 14 (region 40e+07 -100e+07) at a resolution of 40 Kb that in four cell lines (see Supplementary Table 1). In addition to the H1-hESC(stem cellline) cell line, the other three cell lines (A549:lung, G401:kidney, and LNCaP:prostate) are human cancer cell lines (Supplementary Fig 1–2) [37, 38].The change in entropy was confirmed to be significant by computing the p-value Supplementary Fig 5).We observed that despite some changes in the connectivity, the region-specific entropy signal bears striking similarities among the cancer cell lines. In particular, in Fig. 1 B, the entropy is elevated between the 75e + 07 geomice region and 90e + 07 compared to neighboring regions in all cancer cell lines, featuring a similar pattern. This conservation of entropy features is not observed in the stem cell stage (hESC).

We analyze the signal profiles of H3K27ac (acetylation) and H3K27me3 (methylation), key epigenetic markers of chromatin accessibility, together with CTCF binding data (Supplementary Fig 7). The cancer cell line A549 exhibited dense H3K27ac signals, consistent with an open chromatin state, in contrast to the more developmentally poised human embryonic stem cells (hESCs). These patterns align with the entropy landscapes computed using VECTOR, as the A549 cell line displayed pronounced local fluctuations in entropy, suggestive of a more flexible organization of chromatin. In contrast, hESCs showed a smoother, elevated entropy profile, indicating consistent connectivity and a more ordered genome structure. These differences are also evident in the corresponding Hi-C maps, where hESCs display well-defined domain boundaries compared to the more fragmented organization in A549 (Supplementary Fig 1–2).

Our findings highlight the utility of entropy as a quantitative metric for chromatin state, capturing both structural variability and organizational coherence across different cell types. Notably, we observed that genomic regions with higher network connectivity tended to exhibit increased entropy, reinforcing the interpretation of entropy as a readout of 3D genome complexity and plasticity. To explore whether these entropy features may reflect functionally relevant genomic programs, we examined the high-entropy region 75–90 Mb, which showed pronounced variability in A549 cells. Within this region, we identified several genes associated with the Notch signaling pathway along with fatty acid metabolism, two processes known to contribute to cancer progression(Supplementary Fig 6)[36]. While a detailed mechanistic investigation is beyond the scope of this study, we note that several of these genes exhibit cancer-type-specific dysregulation in external datasets, including upregulation of the Notch repressor NUMBin kidney and prostate cancers[39], and altered expression of fatty acid metabolism genes such as ACOT1 and DLST in gastric and neural tumors[40, 41].

Although these observations are preliminary, they suggest that regions marked by elevated entropy may coincide with loci involved in key regulatory and metabolic pathways, potentially reflecting coordinated reprogramming of chromatin architecture and gene function in cancer. These findings support the broader applicability of entropy-based genome analysis for uncovering structural correlates of disease-associated regulatory changes.

We next evaluated the sensitivity of our method in detecting differences in chromatin organization across(NBRs) and within cell lines(BRs). For this, we evaluated the ability of our method to differentiate biological replicates (BRs) from non-biological replicates (NBRs). We found that the entropy profiles of the BRs of the same cell line were more closely clustered along principal component 1 than those of different cell lines (Fig. 1C). This shows that VECTOR is sensitive enough to distinguish between biological replicates. In addition, entropy signals specific to each cell line are conserved, indicating different local chromatin arrangements that regulate gene expression.

#### Sensitivity of VECTOR to stride and resolution changes

We evaluated the robustness of VECTOR to change in stride values and the resolution of the Hi-C contact map. For the first one, we varied the stride sizes during the entropy computation. The stride parameter determines the sampling density from the input contact matrix, which could influence the entropy profile. To evaluate this effect, we systematically modulated the stride size from 2, 5, 10, 15, and 20 and examined the resulting entropy signals. The results presented in (Fig. 2A) indicate that the entropy profiles remain largely consistent across different stride values, with all signals overlapping closely, resembling a moving average. This reflects the robustness of VECTOR against the changes in stride values.

**Fig. 2.**
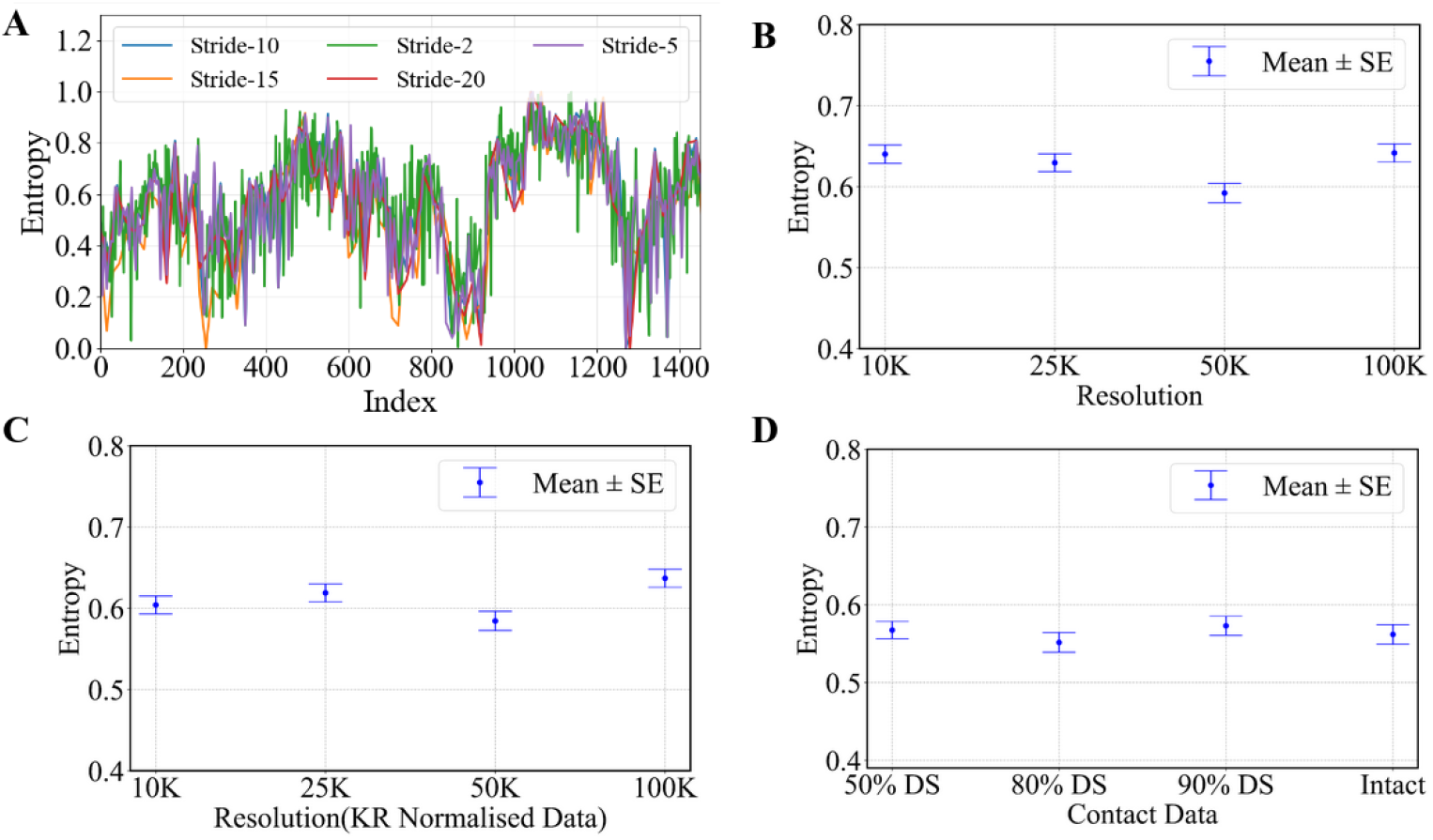
(A) Entropy signals in A549 cell lines estimated using different amounts of input samples, showing overlapping trends similar to a moving average. (B) Impact of resolution changes on entropy signals, revealing minor deviations ranging from 0.58 to 0.6. (C) Entropy signals derived from KR-normalized Hi-C data, exhibiting minor fluctuations while maintaining the overall pattern. (D) Entropy signals in A549 cell lines estimated using varying interaction pairs from Hi-C data (abbreviated as DS: Down-sample), demonstrating that the entropy feature remains largely unaffected by down-sampling .

We further investigated whether the entropy signal captured by VECTOR is generalized and robust across different Hi-C data parameters, such as resolution (binning) and down-sampling. Down-sampling reduces the number of data points while preserve the essential structural features, enabling efficient analysis without a significant loss in interaction information. Consequently, we evaluated the impact of Hi-C map resolution on the magnitude of the entropy signal (seeFig. 2B). Despite changes in the resolution, the mean entropy remains consistent, indicating that VECTOR effectively captures genomic connectivity patterns across varying resolutions. To further ensure that this stability is not influenced by normalization techniques, we evaluated entropy signals in Hi-C data normalized using the Knight-Ruiz (KR) matrix balancing method (Fig. 2C). The results showed that KR normalization maintained a similar entropy pattern, with mean entropy values closely matching those from unnormalized data and only minor fluctuations observed.

To test the robustness of VECTOR under down-sampling conditions, we analyzed the entropy signal at different levels of data reduction: 50% (24M), 80% (39M), and 90% (44M) of the total interactions. As shown in (Fig. 2D), the mean entropy remains consistent across all down-sampling levels, with only minor variations, further demonstrating the method’s reliability. These findings underscore the robustness of VECTOR in estimating genomic entropy across a wide range of analytical conditions. Even with substantial down-sampling (up to 50%, e.g., 24M in A549), varying stride values, and the application of KR normalization, VECTOR consistently distinguishes chromatin architectures. This highlights its reliability and adaptability in capturing biologically meaningful structural differences irrespective of input resolution or preprocessing choices.

#### VECTOR demonstrates superior performance over traditional Spearman and Pearson correlation methods in distinguishing the developmental stages of mouse cells

Due to the limited availability of biological replicates across all samples, we shifted our focus on quantifying intra-cell-type relationships. In particular, we examined whether VECTOR could effectively distinguish different developmental stages within the same cell line, and whether the entropy signal could reveal biologically meaningful changes in genome connectivity associated with the development. We analyzed eight developmental stages in mice: oocyte, PN3 (stage 0; S phase), PN5, early 2-cell (G1), late 2-cell (S–G2), 8-cell, embryo, inner cell mass (ICM), cortex, and mouse embryonic stem cells (mES). Our study focused on architectural details of chromatin in gametes and preimplantation embryos.

Entropy signals were computed for each stage, and to assess their relationships, we calculated pairwise Pearson correlations of these signals and generated an entropy-based similarity matrix (Fig. 3A). This analysis revealed clear separations between developmental stages, suggesting that entropy effectively captures dynamic changes in genome organization. For comparison, we also analyzed contact frequency matrices using both Pearson and Spearman correlations. Pearson correlation values remained uniformly high across all stages (Fig. 3B), leading to an artificial appearance of similarity between distinct stages. Although Spearman correlation (Fig. 3C), which considers rank rather than absolute values, showed reduced similarity, the variation across stages was still minimal, with several stages appearing nearly indistinguishable.

**Fig. 3.**
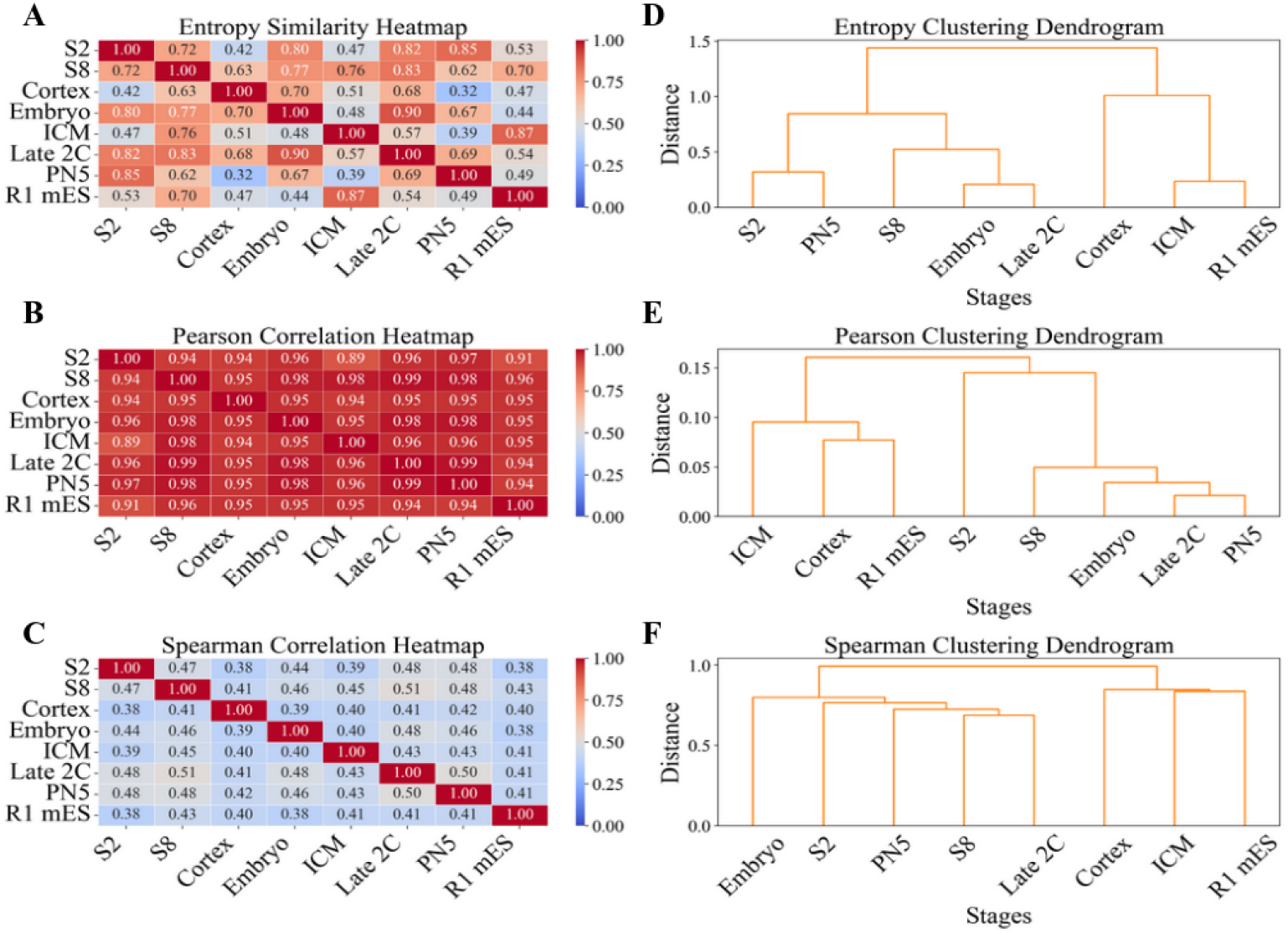
Similarity among Hi-C matrices across developmental stages of the mouse cell line, analyzed using the VECTOR method in comparison with traditional Spearman and Pearson correlation methods. The right panel displays a dendrogram representing similarity relationships.

The similarity matrices are further presented as dendrograms in the right panels of (Fig. 3D–F). The consistent clustering of Cortex, ICM, and R1 mES across all methods reflects their shared mature chromatin architecture, in agreement with observations reported by Du et al. [42]. Beyond these global similarities, the entropy-based dendrogram uniquely resolves local relationships among early developmental stages. Specifically, VECTOR entropy more clearly separates PN3 and early 2-cell stages from later stages such as late 2-cell and 8-cell, capturing the gradual reorganization of chromatin structure during early development. In contrast, dendrograms based on Pearson and Spearman correlations compress many early stages into a single tight cluster, obscuring these biologically meaningful transitions. These local distinctions, evident only in the entropy-based representation, demonstrate VECTOR’s superior sensitivity to fine-grained changes in genome organization. This highlights the utility of entropy signals in defining developmental boundaries and tracking chromatin maturation that is often blurred by traditional contact matrix similarity measures.

To characterize changes in chromatin structure across mouse developmental stages, we computed pairwise differences in entropy, using VECTOR as a measure of chromatin connectivity and structural complexity (Supplementary Fig 8, Table 4). We first tested the normality of entropy distributions using the Shapiro–Wilk test. Since at least one group in each comparison deviated from normality, we used the nonparametric Kruskal-Wallis H test, followed by Dunn’s hoc test with Bonferroni correction for pairwise comparisons. Among the 45 stage-to-stage comparisons, 10 exhibited statistically significant differences in entropy. We observed a progressive increase in entropy from the zygote (stage 0) to later stages, with the 8-cell and Cortex stages showing significantly higher entropy than earlier stages, including stage 0 and stage 2-cell embryos. Specifically, entropy was significantly elevated in stage 8 cells compared to stage 0 (*p <* 0.01) and stage 2-cell (*p <* 0.01), and in Cortex compared to both stage 0 and stage 2. In contrast, a notable decline in entropy was detected in later developmental or pluripotent stages. The ICM showed significantly reduced entropy relative to the 8-cell stage, as did Late 2-cell compared to the 8-cell stage. Similarly, Cortex displayed a significant drop in entropy when compared to ICM and Late 2-cell. R1mES, a pluripotent stem cell line, also exhibited reduced entropy compared to both the 8-cell and Cortex stages.

This non-monotonic pattern, characterized by an initial rise followed by a decline in entropy, suggests dynamic chromatin remodeling during early development. The peak around the 8-cell and Cortex stages is consistent with increased chromatin accessibility and long-range interactions reported in previous Hi-C studies. In particular, the study by Du et al. [42] identified transient long-distance chromatin interactions (2–20 Mb) during the 2-cell to 8-cell stages, which became attenuated in ICM. These features coincide with our entropy peak, supporting the interpretation that higher entropy reflects a structurally dynamic, less compartmentalized chromatin state. The subsequent reduction in entropy in ICM and R1mES corresponds to a shift toward more compact and insulated chromatin, as also evidenced by the emergence of stronger TAD boundaries and reduced distal interactions in Hi-C data.

Overall, our entropy-based analysis provides a quantitative and reproducible framework for detecting chromatin reorganization during early development, capturing known transitions in 3D genome architecture associated with lineage specification and differentiation.

#### Analysis of Chromatin Spatial Organization During Early Embryonic Development

To quantify changes in the spatial organization of chromatin during early embryonic development, we further analyzed Hi-C contact maps by converting them into graphs consisting of 1,000 nodes at a resolution of 50 Kb. The entropy was then calculated as described in previous sections to assess structural variations across different developmental stages. Although previous studies report that topologically associating domains (TADs) emerge as early as the zygote stage, they appear to remain in a primed state, with weak structural consolidation and boundary insulation. We aim to revisit this observation using our entropy-based method, VECTOR, to quantify structural complexity and gain deeper insights into early chromatin organization [42].

To investigate changes in chromatin connectivity across developmental stages, we analyzed local von-Neumann entropy profiles derived from VECTOR. These profiles allowed us to identify genomic regions exhibiting conserved or dynamic structural features during early mouse development. We first compared the entropy values between adjacent developmental stages-specifically, Stage 0 (PN3) versus Stage 2 and Stage 2 versus Stage 8 (Fig. 4). Regions that deviated significantly from the fitted regression line in these scatter plots were indicative of major shifts in chromatin connectivity. Such deviations suggest a change in local structural constraints, potentially reflecting stage-specific chromatin remodeling and gene regulatory activity.To systematically quantify these changes, we calculated the perpendicular distance of each genomic locus from the fitted inter-stage trend line. This distance represents the magnitude of entropy shift between stages. Using these values, we applied K-means clustering to group genomic regions based on their degree of change in connectivity (Fig. 5). Clusters near the regression line (e.g., Cluster 0) represent loci with minimal change in entropy, suggesting similar chromatin structure across developmental transitions. Clusters far from the regression line (e.g., Cluster 3) indicate loci with significant entropy variation, pointing to dynamic chromatin rearrangements. Interestingly, Clusters 2 and 3 (green and yellow) were enriched in genomic regions that later participate in long-range chromatin interactions, supporting the idea that these loci are primed for reorganization during development. To demonstrate the comparative utility of VECTOR, we next identified genomic regions with highly similar entropy profiles across developmental stages at 50 Kb resolution (Fig. 6 and Supplementary Figure 11-13). Similarity was defined based on entropy slope values between stage pairs, with a threshold of slope ≥ 0.85 used to identify conserved regions. Among these, the most similar stage pairs included PN3(stage 0) vs. PN5, Stage 2 vs. Embryo, and Stage 2 vs. PN5 (Fig. 6). For each comparison, the top 10 most similar regions were selected, and associated genes were mapped to these loci. Functional annotation revealed enrichment for genes involved in critical biological processes:The genes located within regions of high entropy similarity are functionally diverse yet closely tied to key developmental processes. Notably, they include patterning and morphogenesis genes such as Hoxd and Evx2; genes essential for tissue-specific structures like Lrp4 and Ptprj; and those involved in neuronal development and function, including Lnpk and Scn7a/Scn9a. Other identified genes contribute to cellular machinery and signaling (Psmd14, Tank, Grb14), as well as immune protection and specialization (Cd59a/b, Olfr1012). These genes are implicated in crucial biological pathways such as limb and digit formation, neural patterning, and immune response, many of which have been previously associated with embryogenesis and organogenesis [43–45]. Together, these findings suggest that genomic regions with conserved entropy profiles across developmental stages may correspond to evolutionarily conserved regulatory hubs that are essential for orchestrating proper developmental progression.

**Fig. 4.**
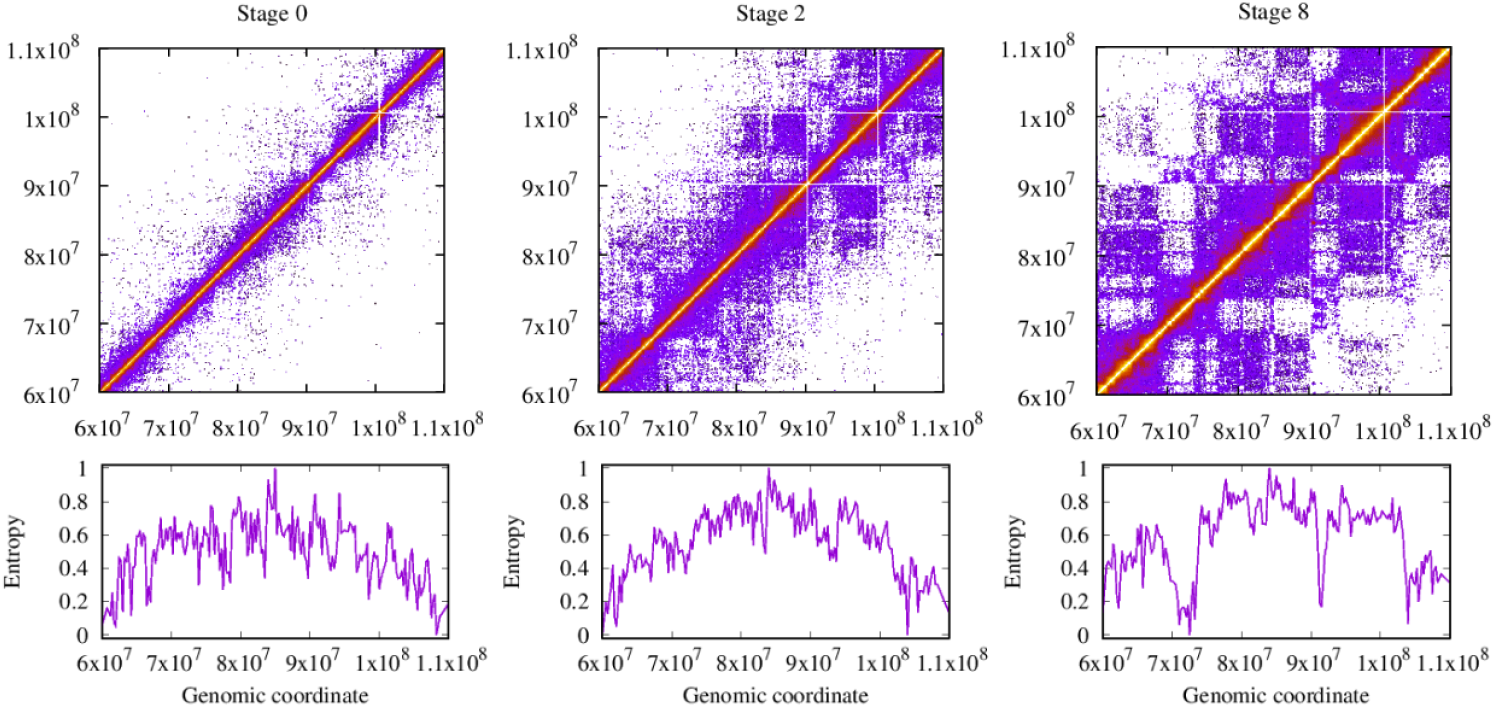
(A) Hi-C contact maps for mouse developmental stages 0(PN3), 2, and 8, with entropy variation shown below.

**Fig. 5.**
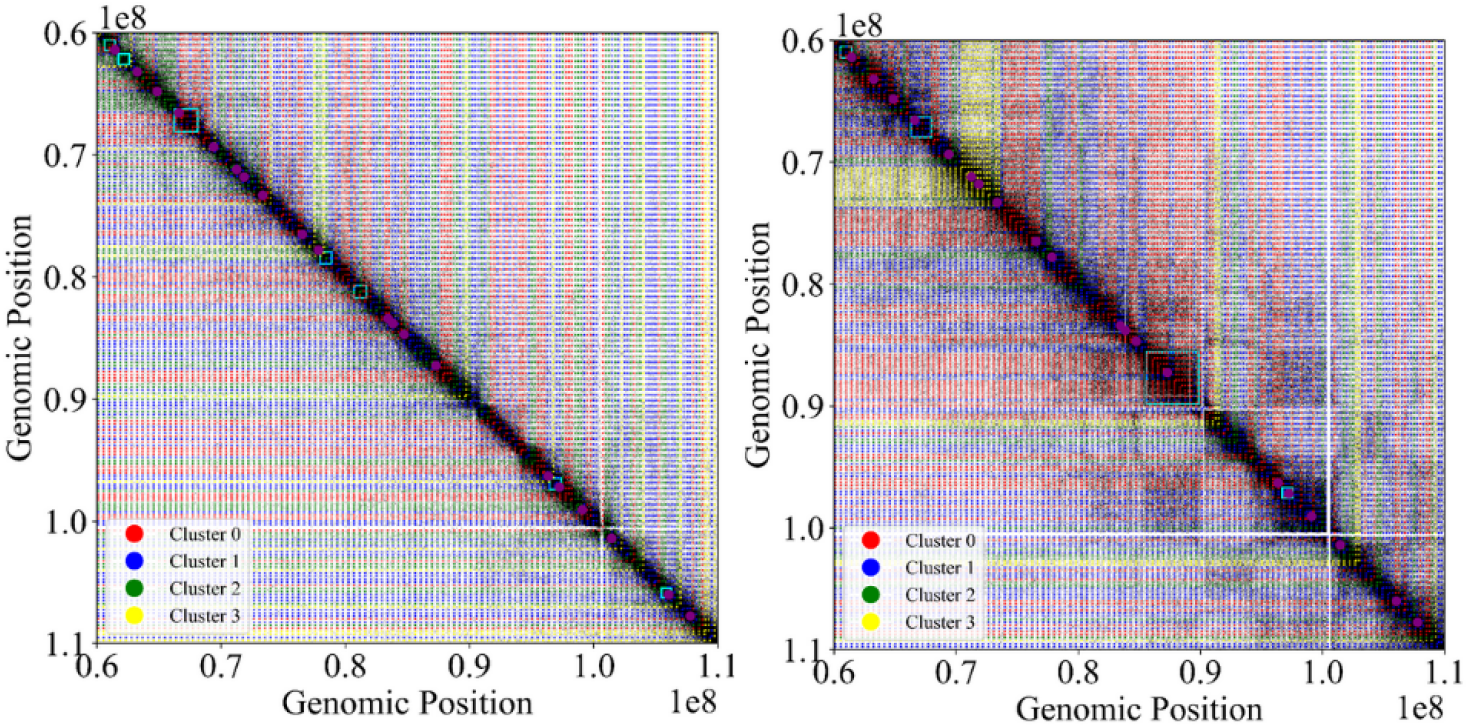
Quantification of entropy variations across consecutive developmental stages, with k-means clustering highlighting regions of similar entropy dynamics. The left panel shows the Hi-C contact map for stage PN3 (stage 0), with clusters derived from entropy changes between stages 0 and 2. The right panel presents the Hi-C map for stage 2, with clusters based on entropy differences between stages 2 and 8. CTCF binding sites (magenta dots) and TAD boundaries (cyan squares) are overlaid to provide structural genomic context.

**Fig. 6.**
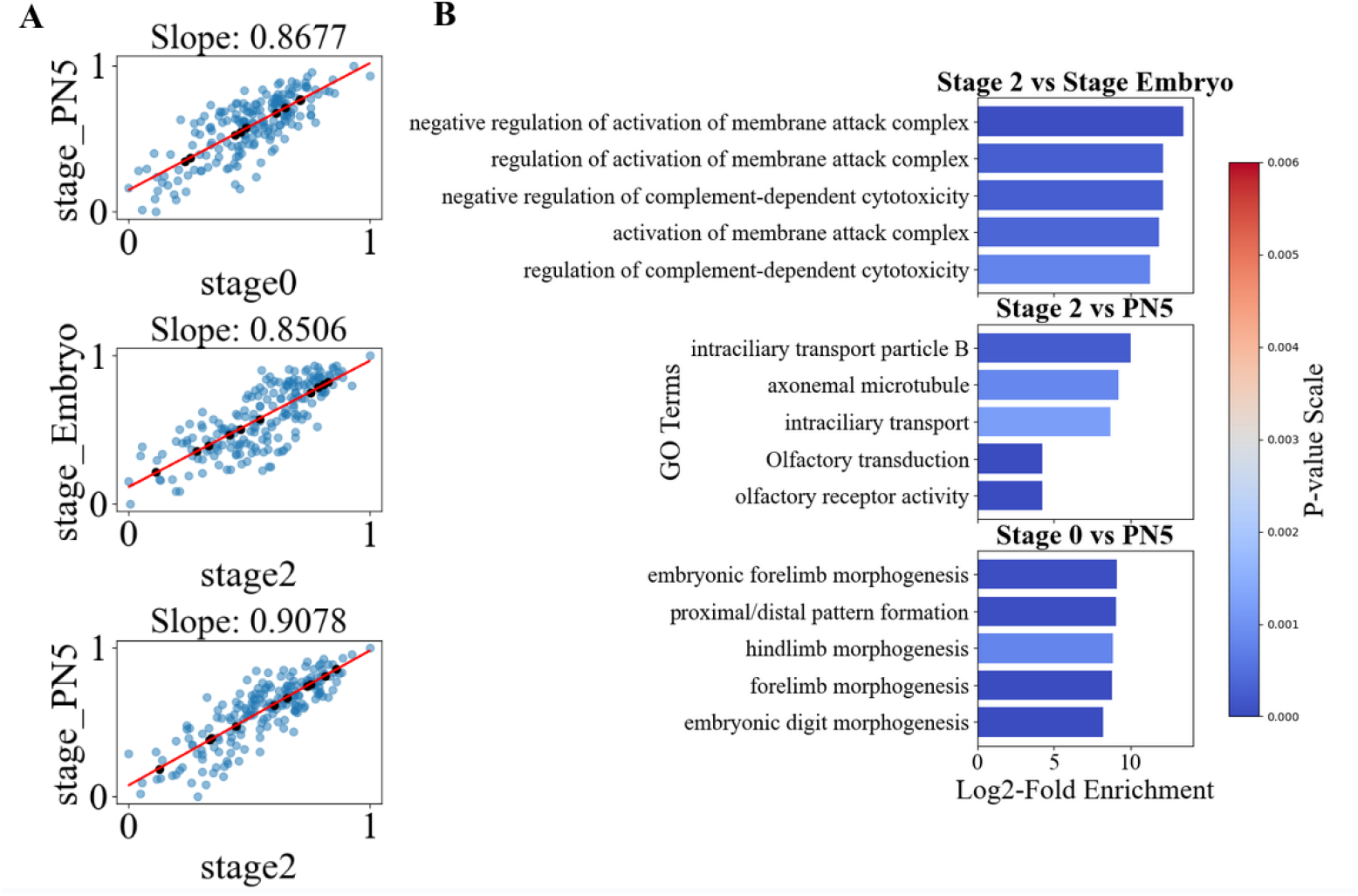
Entropy signals can be utilized to investigate the biological role of regions with similar complexity between two cell stages. (A) Data points represent the entropy values of two compared stages at 50 kb resolution with a stride of 5. The red line represents the fitted regression model, used to identify regions of similar complexity between the stages. The top 10 closest points to the regression line are further analyzed. Black points indicate top 10 data points that has least perpendicular distances from the regression line, representing the regions with the most similar complexity. (B)The top 5 Gene Ontology (GO) terms associated with genes in the most similar regions are shown, ranked by log_2_-fold enrichment. The x-axis represents log_2_-fold enrichment, while the y-axis lists the GO terms. Bars are colored based on p-value significance.

#### VECTOR competes well with other contact matrix similarity methods

VECTOR demonstrates competitive performance compared to other state-of-the-art Hi-C similarity methods. In particular, when benchmarked against HiCRep, VECTOR achieved a wider separation between biological replicate (BR) and non-replicate (NR) similarity scores (see Methods), indicating improved discriminative capability. Specifically, VECTOR yields an average separation margin of *d* = 0.16, which, while lower than ENT3C’s margin (*d* = 0.49), reflects its balanced sensitivity across different interaction scales.

However, it is important to contextualize these results: The reported margin for ENT3C was calculated using small submatrices near the diagonal of the Hi-C contact map, which selectively captures short-range interactions ignoring the long-ranged ones. This performance drop is likely due to the methodological design of ENT3C, which excludes long-range contacts beyond a fixed matrix window size [25]. The assumption potentially misses contacts formed due to long loops. Such omissions may limit its utility in capturing global chromatin organization, where long-range interactions are often functionally relevant. When the analysis scale was expanded to include more than 80%(submatrix dimension = 2050) of the contact matrix, ENT3C performance declined significantly. In some cases, it even failed to correctly distinguish BRs from NBRs, with NBRs exhibiting higher similarity to each other than to their corresponding BRs (Fig. 7). In contrast, VECTOR computes entropy using ego-graphs,each consisting of a node and its immediate neighbors and applies the normalized Laplacian and full-spectrum entropy calculation. This approach captures both short- and long-range interactions, offering a more comprehensive and robust measure of chromatin structure. As a result, VECTOR is particularly well-suited for comparative analyses across large-scale or sparse Hi-C datasets.

**Fig. 7.**
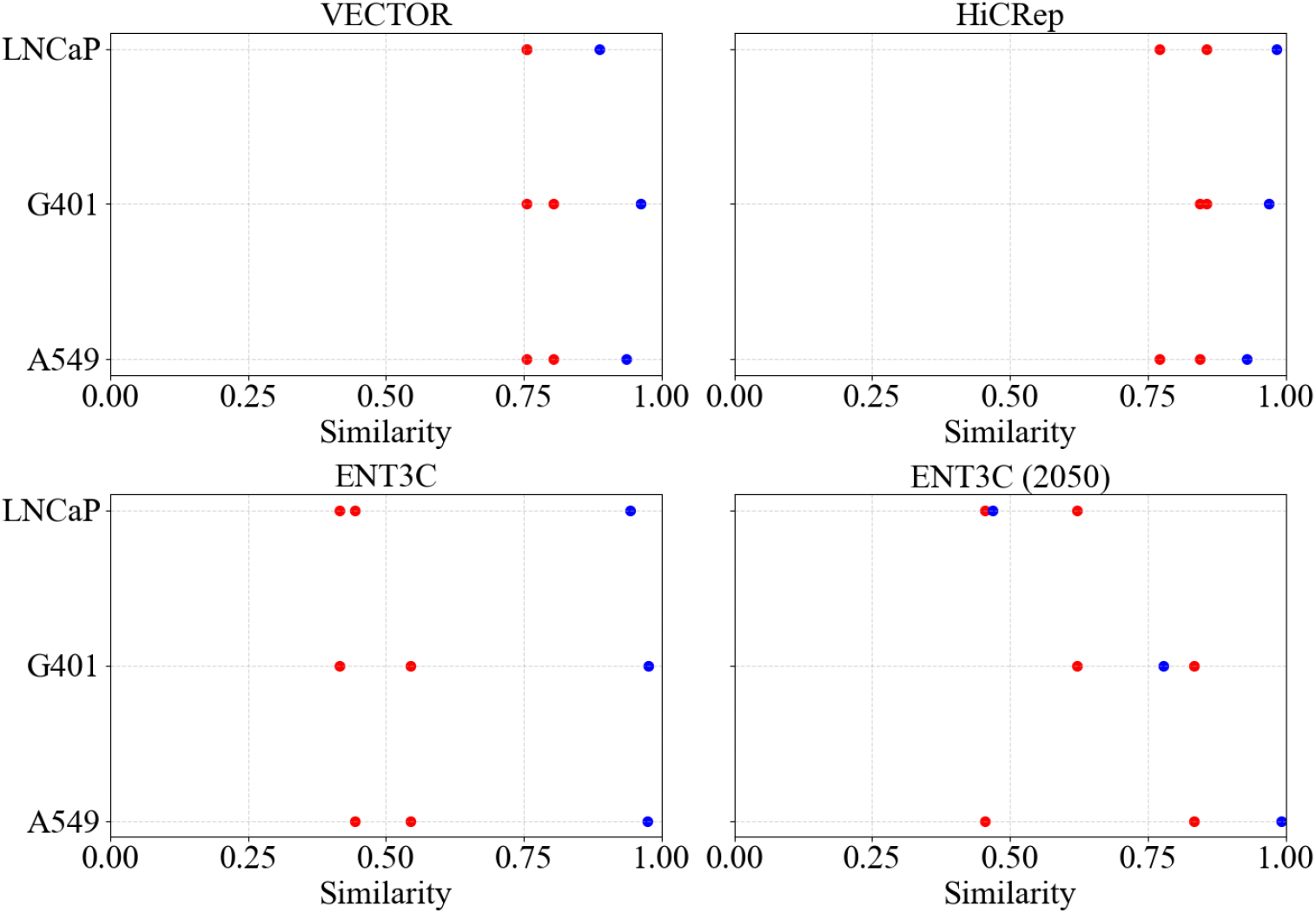
Comparison of VECTOR with other Hi-C similarity quantification methods. The upper panel shows similarity scores from VECTOR (left) and HiCRep(right), while the lower panel presents scores from the von-Neumann entropy-based method and ENT3C. Each dot represents the average similarity score for a given pair of Hi-C matrices. Blue dots indicate biological replicates, and red dots indicate non-biological replicates.

To further assess the applicability of our entropy-based method, we applied VECTOR to sparse single-nucleus Hi-C (snHi-C) data of chromosome 19 from a mouse brain development dataset [46], encompassing postnatal stages P1, P7, and P28. Despite the sparsity and variability inherent in snHi-C data, VECTOR successfully captured structural features that distinguish chromatin organization across developmental timepoints. The PCA plot based on VECTOR-derived entropy signals (Fig. 8)(left panel) shows a clear separation between early (P1 and P7) and late (P28) stages. PC1 explains 33.5% of the variance and strongly differentiates P28 samples (shift along the positive axis), from the initial stages (P1 and P7), which cluster together on the negative side. Although the P1 and P7 samples are not clearly separated from each other, possibly due to minimal changes in chromatin organization, the transition from early to late development is clearly captured by VECTOR. In contrast, PCA using ENT3C features (Fig. 8)(right panel) although explains a higher proportion of variance (48.9% in PC1), but fails to distinguish developmental stages effectively. Samples from different timepoints (e.g., P1-2 and P28-2) overlap substantially in the PC space, and stage-specific clustering is less evident. The findings clearly demonstrate that VECTOR is more effective than ENT3C in detecting large-scale chromatin reorganization during developmental transitions, particularly in distinguishing early postnatal stages from later ones. Its ability to extract consistent and biologically meaningful patterns, even from sparse Hi-C datasets, underscores its potential for analyzing global structural changes across cell states or time points.

**Fig. 8.**
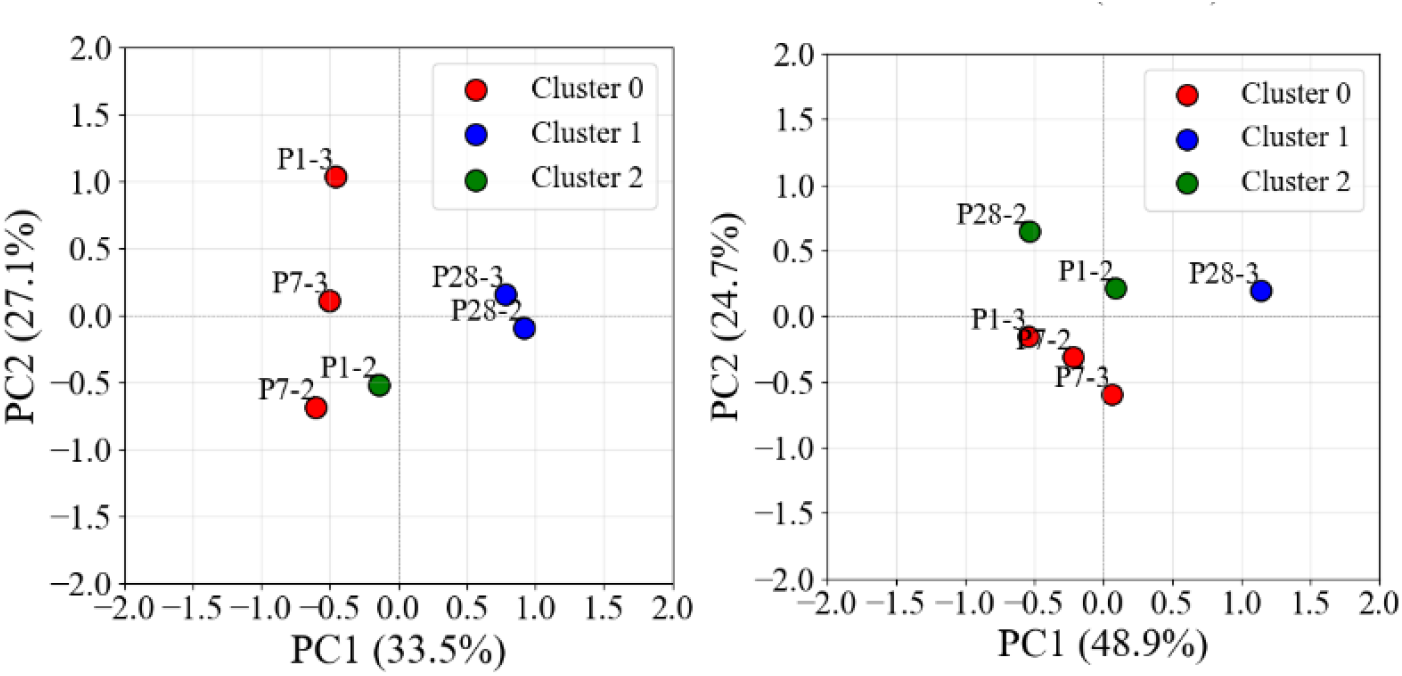
Principal component analysis (PCA) of entropy features from single cell Hi-C data across developmental stages. The left panel shows PCA based on VECTOR-derived entropy signals, where PC1 (33.5% variance explained) clearly separates early (P1, P7) and late (P28) developmental stages. Clusters are compact and well-separated, reflecting stage-specific chromatin organization. In contrast, the right panel shows PCA using ENT3C features, where PC1 explains more variance (48.9%) but shows less distinct stage separation, with noticeable overlap between samples from different stages

## Discussions and Conclusions

In this study, we introduced VECTOR, a novel computational approach leveraging Von Neumann entropy to quantify structural variations in chromatin organization from Hi-C contact maps. Our results demonstrate that VECTOR effectively captures the complexity of 3D genome architecture and provides a robust measure of chromatin connectivity across different biological conditions, including various cell lines and developmental stages.

By applying VECTOR to human chromosome 14 across multiple cell lines, we observed distinct entropy patterns reflecting variations in chromatin structure. Notably, entropy profiles among cancer cell lines exhibited significant conservation, whereas the stem cell line (hESC) displayed greater structural variability.Detailed comparison between A549 and hESC cells (Supplementary Fig 7) reveals marked differences in chromatin organization. The entropy patterns reinforce our results, with A549 exhibiting greater disorder and variability, and hESCs maintaining a more stable and organized chromatin architecture, consistent with Hi-C and epigenetic profiles (Supplementary Fig 1–2).This suggests that VECTOR entropy signals can serve as reliable indicators of chromatin connectivity changes associated with cellular differentiation and disease states. Additionally, VECTOR proved its sensitivity in distinguishing biological replicates from non-biological replicates, reinforcing its reproducibility and robustness in Hi-C data analysis similar to other reported startes of arts such as ENT3C [25].

Furthermore, our systematic analysis of stride and resolution dependencies of VECTOR confirmed the stability of VECTOR-derived entropy profiles. Across a range of stride values and resolutions, the entropy signals remained consistent, demonstrating the method’s robustness against minor parameter variations. This stability ensures that VECTOR can be applied to Hi-C datasets of varying resolutions without introducing significant distortions in the detection of chromatin structural patterns.

Further analysis on mouse cell-lines demonstrated the superior performance of VECTOR over traditional correlation-based methods such as Pearson and Spearman correlation in distinguishing distinct embryonic developmental stages. While conventional methods exhibited limitations in resolving chromatin structural differences between closely related stages, VECTOR successfully identified key genomic regions undergoing dynamic reorganization. Our findings highlight the utility of entropy-based metrics in capturing meaningful structural transitions that are essential for understanding genome regulation during development.

Additionally, the application of VECTOR to early embryonic development revealed genomic regions with distinct connectivity shifts between successive stages. By quantifying entropy variations and clustering regions based on connectivity changes, we identified structural transitions that correlate with key regulatory elements such as TADs and CTCF binding sites. These observations suggest that entropy-derived measures provide insights into chromatin remodeling events, shedding light on the mechanistic basis of genome organization changes during cellular differentiation.

A key determinant of the quality of the entropy signal lies in the choice of the density matrix used in the spectral entropy formulation [26]. To systematically investigate this, we compared entropy signals derived from various Laplacian-based constructions (see Supplementary Fig. 9).We found that the entropy signals derived from the standard and normalized Laplacians exhibit broadly similar trends. In contrast, the exponential Laplacian—while theoretically satisfying the subadditivity property—produces higher-frequency fluctuations and exhibits sensitivity to sparse matrix noise, which can obscure biologically meaningful patterns in Hi-C data. Based on these observations, we adopted the normalized Laplacian in VECTOR as the default because it balances sensitivity with robustness across heterogeneous datasets, preserves interpretability, and avoids scale-dependent artifacts. This choice supports consistent biological conclusions while aligning with recent literature advocating for the normalized Laplacian in spectral entropy analyses.

To assess VECTOR’s effectiveness, we benchmarked its performance against two leading Hi-C comparison tools: ENT3C [25] and HiCRep [10]. VECTOR consistently outperformed both methods in distinguishing non-biological replicates (NBRs), demonstrating heightened sensitivity to biologically meaningful differences in chromatin conformation. Notably, it retained this discriminative power even in sparse single-cell datasets, revealing reproducible entropy signatures across biological replicates (BRs) and capturing distinct structural patterns between unrelated samples. These findings underscore VECTOR’s robustness and versatility across a range of Hi-C data types, including low-coverage, single-nucleus, and developmental time-series datasets, where conventional methods often fail. Using the Laplacian normalized graph to integrate both short- and long-range interactions, VECTOR produces a entropy measure that reflects the global topological organization of the genome. This holistic perspective enables more reliable comparisons between heterogeneous cellular states, developmental stages, or experimental conditions.

Unlike ENT3C, which is constrained by its emphasis on local, short-range interactions, VECTOR is specifically designed to capture the multiscale nature of chromatin architecture. Although interpreting entropy changes in terms of specific biological processes remains challenging, VECTOR effectively quantifies structural similarity and reveals meaningful patterns of chromatin reorganization. Its capacity to extract consistent, interpretable features from even sparse contact maps positions it as a valuable tool for investigating dynamic genome organization in both basic and translational research contexts.

In line with previous findings that TAD-like domains (TLDs) persist at the single-cell level and correspond well with ensemble Hi-C maps[15, 47], we assessed whether entropy profiles generated by VECTOR could similarly reflect conserved chromatin architecture. Despite the sparsity of single-cell Hi-C data, the entropy profiles derived from pooled and ensemble datasets [48] exhibit strong global similarity, with rolling-average Pearson correlations exceeding 0.85. This suggests that higher-order chromatin organization is largely maintained even when limited cell numbers are used (Supplementary Fig 10).

Importantly, prominent entropy peaks and valleys closely align with domain boundaries identified by deDoc2[47], reinforcing the utility of entropy as a robust and interpretable proxy for 3D genome structure. While exact local correspondence between entropy dips of pooled and ensemble is not always observed, likely due to the stochastic nature of single-cell contact patterns. However, the overall agreement highlights VECTOR’s ability to capture consistent chromatin features under experimentally constrained conditions. .

To summarize, our study establishes VECTOR as a powerful and computationally efficient tool for characterizing 3D genome organization. Its ability to differentiate structural variations across biological conditions, its robustness to parameter changes, and its superior performance in developmental stage differentiation make it a valuable addition to the existing repertoire of Hi-C analysis methods. Future applications of VECTOR could extend to diverse biological systems and multi-omics data integration. Further refinement of entropy-based approaches, in conjunction with complementary genomic assays, will enable a deeper understanding of genome architecture and its role in gene regulation.

## 4 Acknowledgements

We gratefully acknowledge the financial support from DST India (CRG/2023/000636), DBT India (BT/PR46247/BID/7/1015/2023) and DBT CoE research grant. A.B gratefully acknowledges support from the Alexandar von-Humboldt Foundation, Germany. D.H gratefully acknowledges the support from Deutsche Forschungsgemeinschaft (DFG, German Research Foundation) under Germany’s Excellence Strategy EXC 2181/1 - 390900948 (the Heidelberg STRUCTURES Excellence Cluster). K. P. K acknowledges the financial support from the DST-SERB project (CRG/2023/000636).

## 5 Data availability

VECTOR is publicly available in Python on GitHub(https://github.com/Kavana-Priya/VECTOR/).

