## Supplementary Information for "VECTOR: A Framework to Identify and Quantify Structural Changes in Chromatin using Hi-C Data"

Kavana Priyadarshini Keshava[1], Dieter W. Heermann [2], Arnab Bhattacherjee[1,2*]

[1*]School of Computational and Integrative Sciences, Jawaharlal Nehru University, New Delhi, 110067, New Delhi, India.
[2]Institute for Theoretical Physics, Heidelberg University, Philosophenweg 19,69120 Heidelberg, Germany.

Contributing authors:;
;

### HiC Data Retrieval

For human cell lines, paired-end sequencing data in BAM format were first converted into the .pairs format using the Pairix pipeline. This involved extracting read-pair information, filtering for valid Hi-C contacts, and storing them in a standardized tab-delimited format that includes chromosome coordinates, strands, and mapping quality for each read.The resulting .pairs files were then used as input to the Juicer Tools pipeline to generate .hic files at 40 kb resolution. Juicer Tools processed these contact pairs to construct normalized Hi-C contact matrices, incorporating binning, filtering, and matrix balancing (e.g., via KR or NONE). These .hic files served as the basis for all downstream analyses, including entropy-based contact map assessments and visualization.

The mouse .hic files used in this study were obtained from Du et al. (2017), and were sourced from the 4D Nucleome database (https://data.4dnucleome.org/). These datasets were originally processed and stored in the .hic format, and analyzed at a 50 kb resolution using Knight-Ruiz (KR) normalization to ensure balanced contact matrices.For compatibility with other comparative entropy tools, the .hic files were converted to the .cool format using the hic2cool utility. This conversion enabled efficient manipulation, visualization, and integration with tools that operate natively in the Cooler format, facilitating standardized analysis across datasets.

We investigated whether sparse single-nucleus Hi-C (snHi-C) data could be effectively distinguished using the entropy-based method. To this end, we utilized snHi-C datasets from mouse brain development as published by Tan et al. (2021) . Specifically, we focused our analysis on chromosome 19 (Chr19) to assess the feasibility of entropy-based differentiation in sparse contact maps.

**Table 1** Human Cell Line Hi-C Files used in this study, along with their biological replicates and accession IDs for BAM files.

| Cell Line | Biological Replicate (BR) | BAM Accession |
|---|---|---|
| A549 | 1 | ENCFF867DCM |
| A549 | 2 | ENCFF532XBC |
| G401 | 1 | 4DNFIU832CGU |
| G401 | 2 | 4DNFIQZ5VU58 |
| hESc | 1 | 4DNFI5M6BXPO |
| hESc | 2 | 4DNFIA3TKHP5 |
| LNCaP | 1 | ENCFF977XHB |
| LNCaP | 2 | ENCFF204XII |

**Table 2** Mouse Cell Line Hi-C files sourced from the 4D Nucleome database and used in this study.

| Cell Stage | Hi-C File |
|---|---|
| 8-cell | 4DNFIFA89L5B.hic |
| Late 2-cell | 4DNFICXCFGEI.hic |
| Early 2-cell | 4DNFIK4CECUH.hic |
| PN5 | 4DNFI1EYIGOC.hic |
| PN3 | 4DNFI7TLEWUI.hic |
| ICM | 4DNFIK5HY1GP.hic |
| Embryo | 4DNFIX4DLXSE.hic |
| R1 mES | 4DNFI5IAH9H1.hic |
| Cortex | 4DNFII3JV8I1.hic |

**Table 3** Mouse cortex Hi-C files from GEO, corresponding to postnatal day 1 (P1), day 7 (P7), and day 28 (P28), used in this study.

| Developmental Stage | Hi-C File |
|---|---|
| P1 cortex, cb 002 | GSM4382150_cortex-p001-cb_002.contacts.hic |
| P1 cortex, cb 003 | GSM4382151_cortex-p001-cb_003.contacts.hic |
| P7 cortex, cb 002 | GSM4382438_cortex-p007-cb_002.contacts.hic |
| P7 cortex, cb 003 | GSM4382439_cortex-p007-cb_003.contacts.hic |
| P28 cortex, cb 002 | GSM4382630_cortex-p028-cb_002.contacts.hic |
| P28 cortex, cb 003 | GSM4382631_cortex-p028-cb_003.contacts.hic |

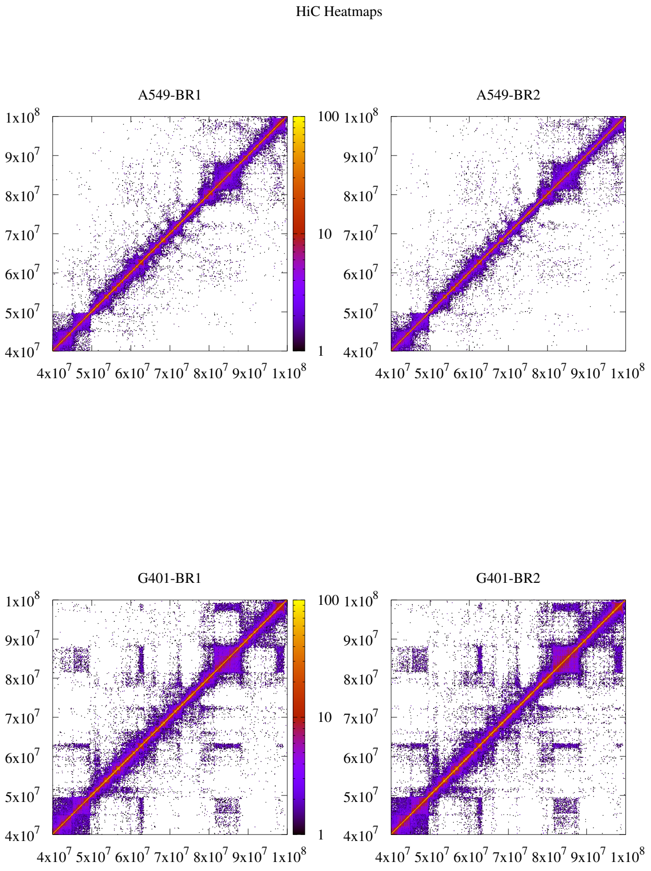


**Fig. 1** Hi-C contact maps of chromosome 14 from human cell lines (A549 and G401) at 40 kb resolution, shown for biological replicates.

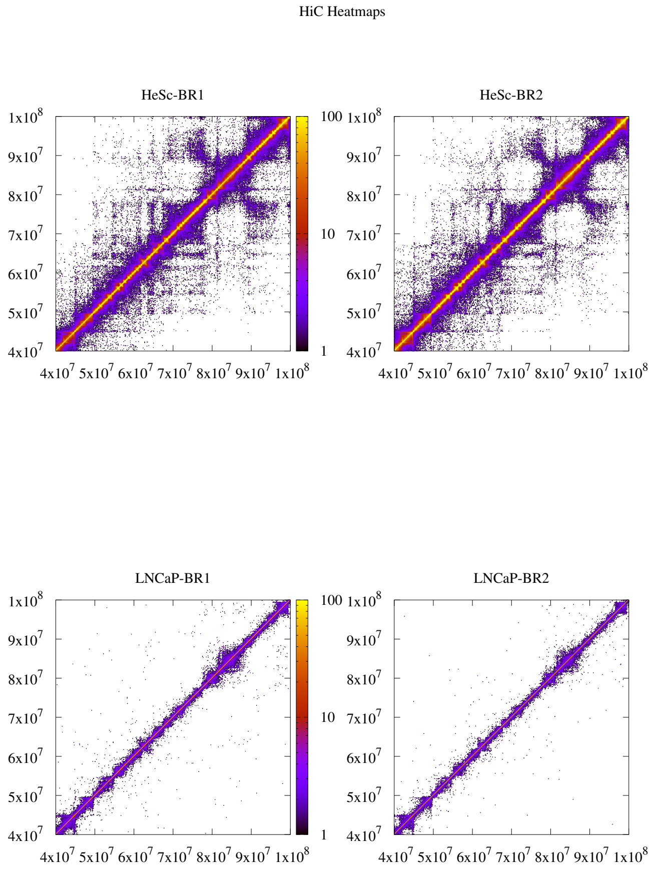


**Fig. 2** Hi-C contact maps of chromosome 14 from human cell lines (hESc and LNCaP) at 40 kb resolution, shown for biological replicates.

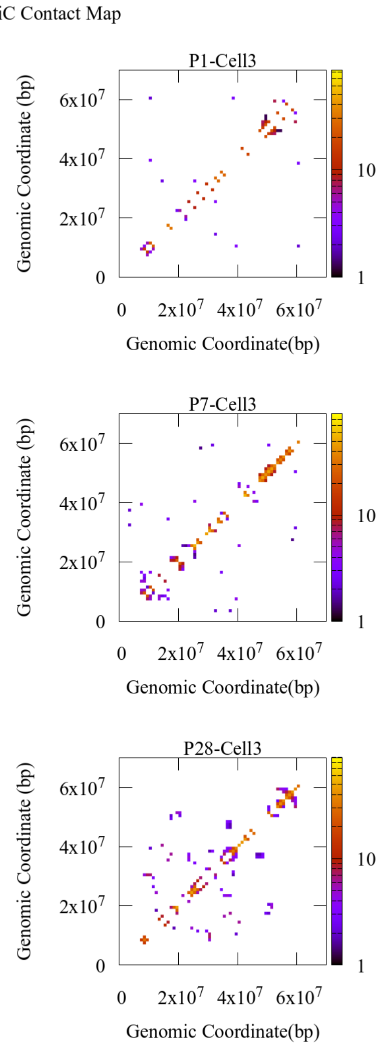


**Fig. 3** Contact maps of chromosome 19 from mouse cortex single cell hic at 1Mb resolution.

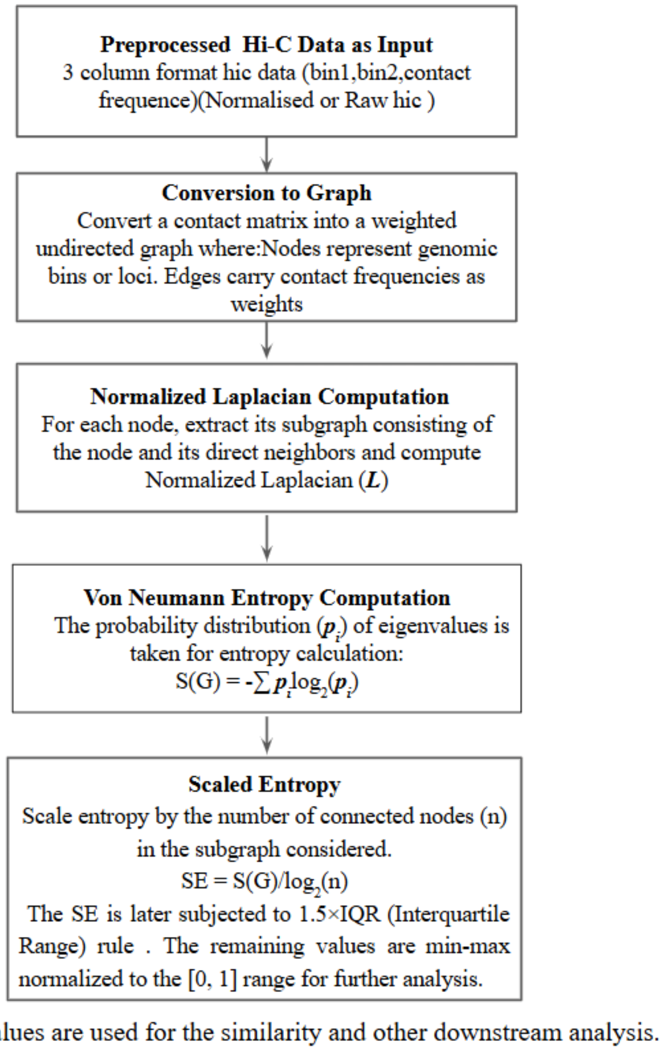


**Fig. 4** VECTOR Workflow.

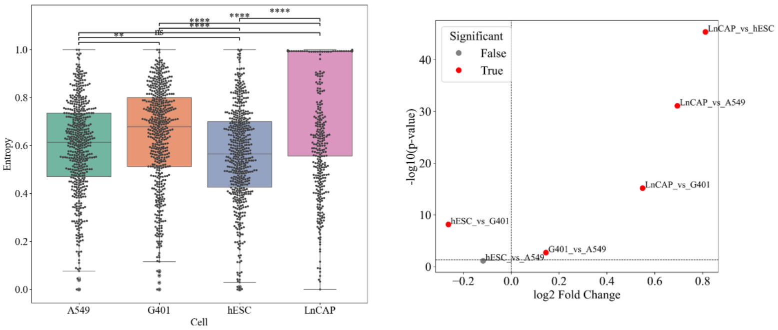


**Fig. 5** The left panel compares entropy across four cell types: A549, G401, hESC, and LNCaP. The LNCaP cell line shows the highest entropy, with several significant differences observed, particularly when compared to hESC and A549. The right panel confirms these findings by highlighting strong statistical significance (red points) in comparisons such as LNCaP vs. hESC, LNCaP vs. A549, and LNCaP vs. G401, with large $\log_2$ fold changes and high $-\log_{10}$ p-values. Non-significant comparisons (e.g., hESC vs. A549 and G401 vs. A549) are shown in grey, suggesting similar entropy distributions among these cell types.

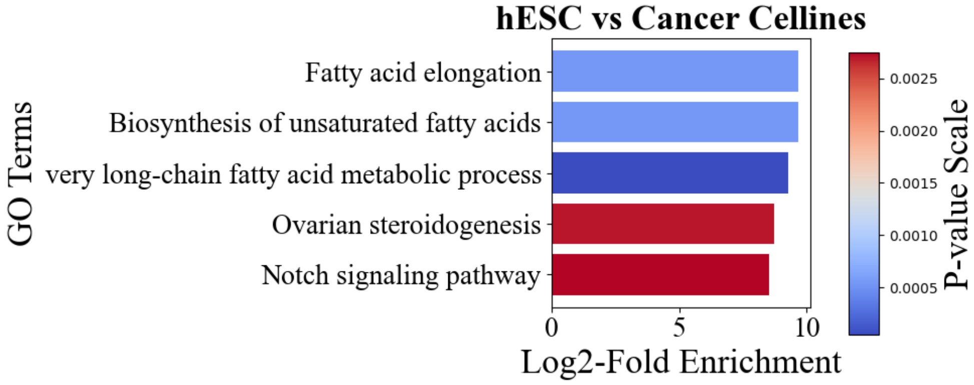


**Fig. 6** The Gene Ontology (GO) terms associated with genes in the most similar regions are shown, ranked by log2-fold enrichment. The x-axis represents log2-fold enrichment, while the y-axis lists the GO terms. Bars are colored based on p-value significance.

### Data Integration

For the human embryonic stem cell line (hESC), we used the following signal tracks: CTCF (`ENCFF269OPL.bigWig`), H3K27ac (`ENCFF103PND.bigWig`), and H3K27me3 (`ENCFF927FVH.bigWig`). For the A549 cell line, the corresponding tracks included CTCF (`ENCFF029NYA.bigWig`), H3K27ac (`ENCFF730VLE.bigWig`) , and H3K27me3 (`ENCFF467XRX.bigWig`).

These tracks were processed and normalized to generate co-plots with computed chromatin entropy profiles, enabling integrative visualization of regulatory features in relation to global chromatin complexity.

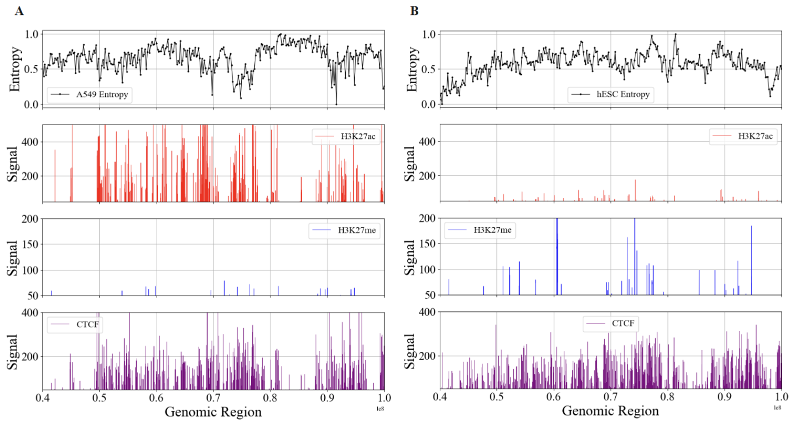


**Fig. 7** Comparison of chromatin entropy and epigenetic landscape across a representative genomic region in A549 and hESC cells. (A) Tracks for A549 cells showing (top to bottom): normalized entropy profile (black), H3K27ac signal (red, active enhancer mark), H3K27me3 signal (blue, repressive histone mark), and CTCF signal (purple, architectural protein). (B) Corresponding tracks for human embryonic stem cells (hESC) across the same genomic region. The x-axis represents genomic coordinates (in base pairs), and the y-axes represent normalized entropy (top panels) or ChIP-seq signal intensity for each epigenetic marker (lower panels).

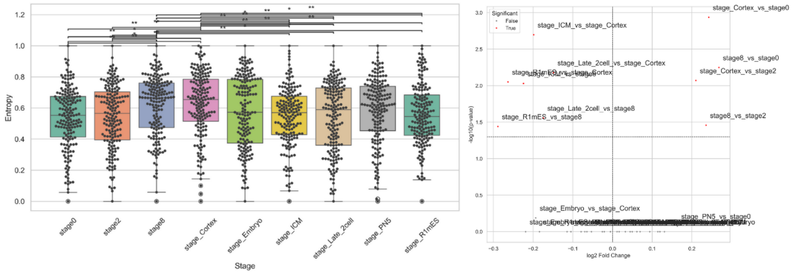


**Fig. 8** In the figure, the box plot (left panel) displays the distribution of entropy across nine distinct embryonic stages: `stage0`, `stage2`, `stage8`, `stage_Cortex`, `stage_Embryo`, `stage_ICM`, `stage_Late_2cell`, `stage_PN5`, and `stage_R1mES`. Significant pairwise differences are indicated by asterisks above the bars, with several comparisons showing high statistical significance (e.g., `stage_Cortex` vs. other stages). (right panel) Plot shows the $\log_2$ fold change in entropy versus the $-\log_{10}$ p-values for each pairwise comparison. Red dots highlight statistically significant comparisons, such as `stage_Cortex` vs. `stage0`, `stage8` vs. `stage0`, and `stage_ICM` vs. `stage_Cortex`, indicating notable differences in entropy between specific stages.

**Table 4** Significant pairwise comparisons of entropy between developmental stages. Positive log2FC indicates increased entropy in the first stage of the comparison.

| Comparison | log2FC | p-value | Direction |
|---|---|---|---|
| stage8 vs stage0 | 0.2682 | 0.0056 | Up |
| stage Cortex vs stage0 | 0.2426 | 0.0012 | Up |
| stage8 vs stage2 | 0.2357 | 0.0348 | Up |
| stage Cortex vs stage2 | 0.2102 | 0.0085 | Up |
| stage ICM vs stage8 | -0.2242 | 0.0093 | Down |
| stage Late_2cell vs stage8 | -0.1743 | 0.0278 | Down |
| stage R1mES vs stage8 | -0.2888 | 0.0362 | Down |
| stage ICM vs stage Cortex | -0.1987 | 0.0020 | Down |
| stage Late_2cell vs Cortex | -0.1487 | 0.0066 | Down |
| stage R1mES vs Cortex | -0.2632 | 0.0089 | Down |

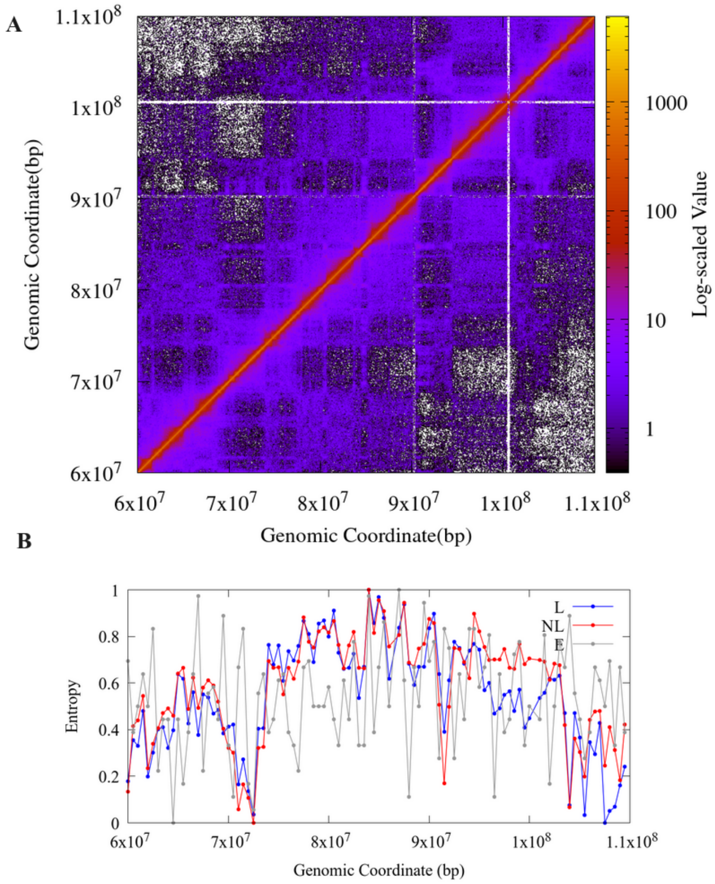


**Fig. 9** The plot shows the scaled von Neumann entropy profiles computed from three types of graph Laplacian-based density matrices applied to mouse chromosome data.(A) Hi-C contact maps of chromosome 2 (from 60 to 110 Mb) in mouse embryonic cells at stage 8, analyzed at 50 kb resolution.(B) Entropy versus genomic coordinates. The blue curve (L) represents entropy derived from the standard Laplacian matrix, the red curve (NL) corresponds to the normalized Laplacian, and the grey curve (E) represents entropy calculated using the exponential of the Laplacian as the density matrix. These entropy profiles reveal differences in structural complexity and information distribution across the chromosome region, depending on the spectral form of the Laplacian used.

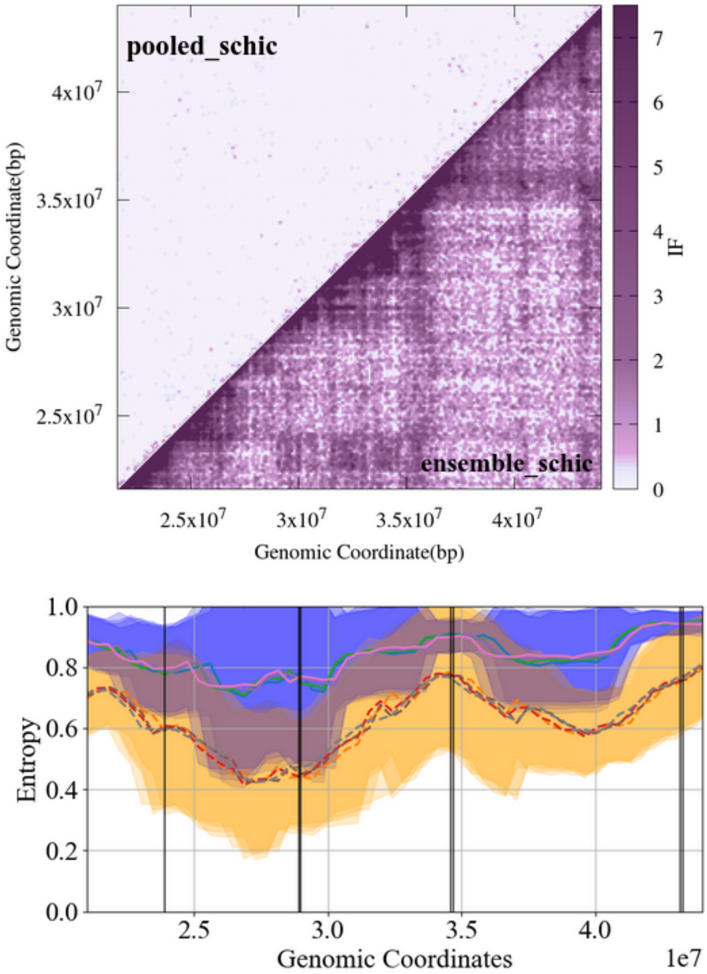


**Fig. 10** (A) Hi-C contact maps for chromosome 11 (85 kb resolution) from Nagano et al., showing pooled single-cell Hi-C (10 cells) and ensemble Hi-C (bulk, 60 cells). (B) Scaled entropy profiles computed using VECTOR for pooled single-cell Hi-C data (solid line) and ensemble Hi-C data (dashed line). TAD-like domain (TLD) boundaries identified by deDoc2 are overlaid. Entropy profiles were smoothed using a rolling window (window size 10–13), yielding a Pearson correlation ranging from 0.8573 to 0.9107 between pooled and ensemble profiles, indicating strong concordance in large-scale chromatin organization.

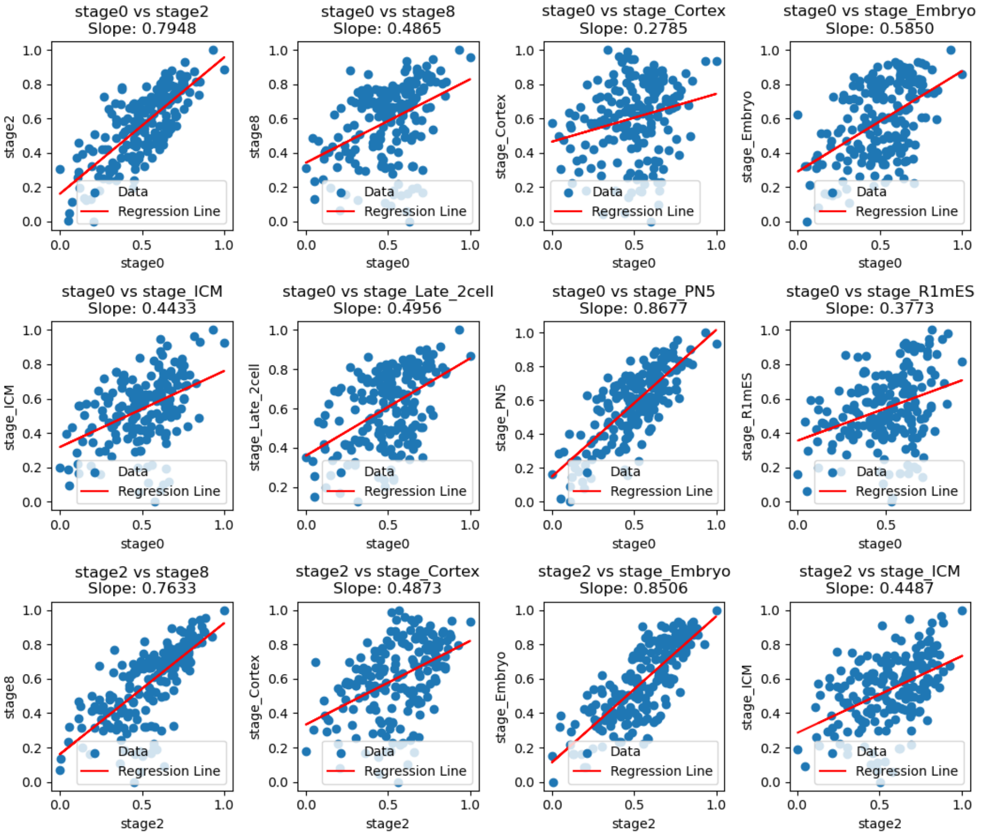


**Fig. 11** Data points represent the entropy values of two compared stages at 50 kb resolution with a stride of 5. The red line represents the fitted regression model, used to identify regions of similar complexity between the stages.

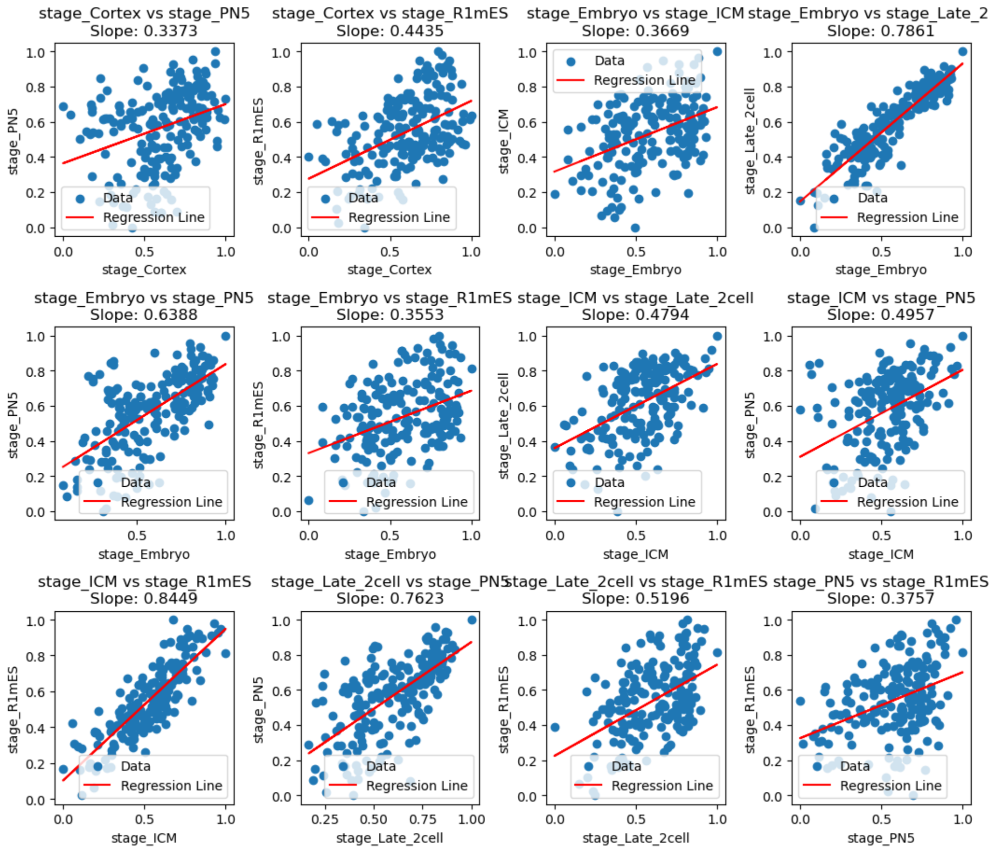


**Fig. 12** Data points represent the entropy values of two compared stages at 50 kb resolution with a stride of 5. The red line represents the fitted regression model, used to identify regions of similar complexity between the stages.

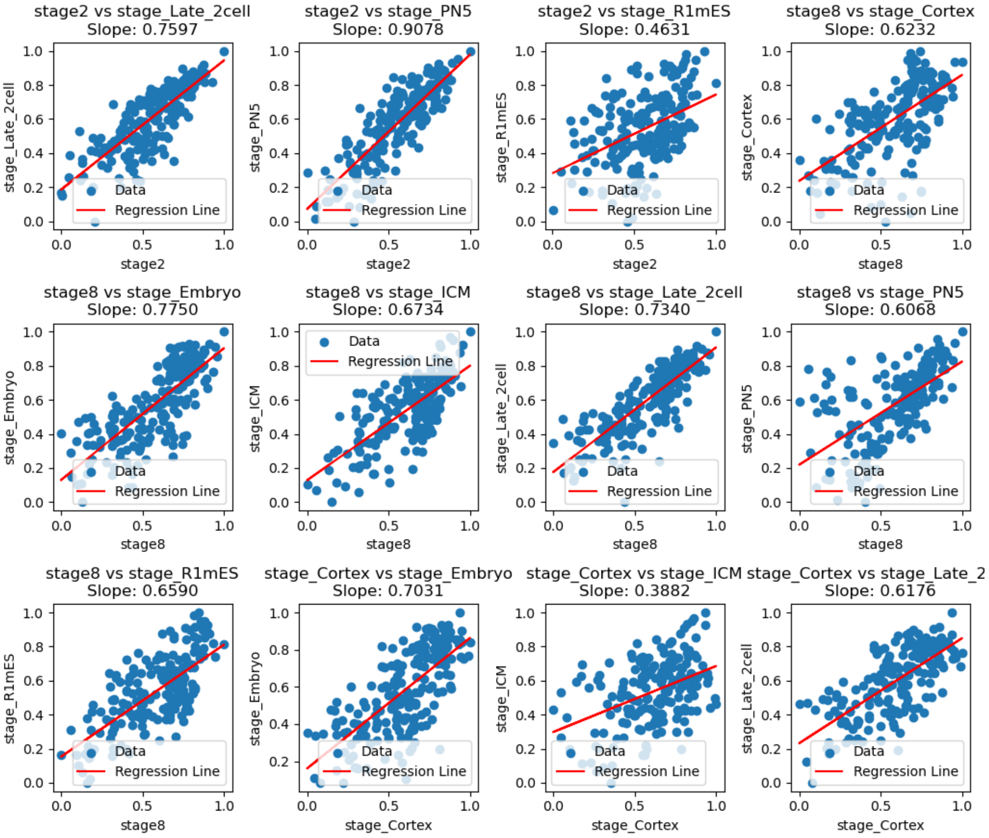


**Fig. 13** Data points represent the entropy values of two compared stages at 50 kb resolution with a stride of 5. The red line represents the fitted regression model, used to identify regions of similar complexity between the stages.
